# Genetic risk score for Alzheimer’s disease is associated with poor hearing

**DOI:** 10.1101/556506

**Authors:** Willa D. Brenowitz, Teresa J. Filshtein, Kristine Yaffe, Stefan Walter, Thomas J. Hoffmann, Eric Jorgenson, Rachel A. Whitmer, M. Maria Glymour

## Abstract

**Introduction:** Incipient dementia or shared genetics may partly explain the association between hearing loss and dementia. We evaluated whether genetic variants known to increase Alzheimer’s disease (AD) also influence hearing difficulty.

**Methods:** UK Biobank participants aged 56+ with Caucasian genetic ancestry self-reported difficulty hearing and hearing with background noise (n=244,915) and underwent objectively measured hearing assessments (n=80,074). Poor objective hearing was defined as >-5.5 dB speech reception threshold on a Digit Triplet Test. We evaluated whether an AD genetic risk score (AD-GRS; range −1.2 to 1.9), the weighted sum of 23 previously identified AD-related polymorphisms, predicted objective or self-reported poor hearing, using age, sex, and genetic ancestry adjusted logistic regression models.

**Results:** Higher AD-GRS predicted objectively measured poor hearing (OR=1.06; 95% CI: 1.01, 1.11) and self-reported problems hearing with background noise (OR=1.03; 95% CI: 1.00, 1.05).

**Discussion:** Using novel methods, we found evidence that AD genetic risk influences hearing loss.

## INTRODUCTION

Moderate to severe hearing loss is estimated to affect up to 40% of adults aged 65 and older and up to 90% of those aged 90+ [1–3]. Several intriguing studies have found hearing impairment is associated with risk of dementia and cognitive decline independent of age and common comorbidities [4–7]. Individuals with poor hearing, whether measured by self-report[8], speech recognition[9], or pure audiometric tone [5,6,10] have elevated risk of adverse cognitive outcomes. Because hearing loss may be preventable or mitigated with the use of hearing aids, it has been suggested that prevention or treatment of hearing loss may reduce dementia risk. Hearing loss was highlighted as one of the most important potentially modifiable risk factors for dementia in a recent Lancet commission on dementia prevention, with a population attributable risk higher than that of the APOE ε4 allele (9% vs 7%) [11]. However, whether hearing impairment causes increased risk of dementia is still debated.

Multiple plausible mechanisms have been posited to explain how hearing loss might cause dementia, including the influence of hearing loss on social isolation or reduced cognitive reserve [12,13]. On the other hand, sensory and cognitive impairments are both strongly associated with age, tend to have a gradual onset occurring across many years, and may have shared etiologies [6,12]. Underlying neurodegeneration or early cognitive changes associated with Alzheimer’s disease (AD) and related dementias may influence hearing testing or processing abilities [14,15]. Other shared diseases, such as vascular or metabolic disease, may affect hearing abilities [16–18] as well as promote cognitive decline [19–21]. Thus, it is plausible that the association between hearing impairment and dementia found in observational studies may be explained by confounding by a shared etiology and/or reverse causation from early neurodegenerative stages of the AD disease process to hearing loss.

Disentangling these alternative explanations for the association between hearing loss and dementia is critical to evaluate the potential for interventions on hearing to reduce dementia burden. If either confounding or reverse causation explain the association, interventions on hearing loss are not likely to reduce dementia risk. Given the long, insidious development of both conditions, conventional regression models cannot establish the temporal order or rule out non-causal explanations for the association. A modified Mendelian randomization approach, based on using genetic risk factors known to increase dementia risk [22,23], can help establish temporal order and evaluate whether underlying dementia-related diseases or shared etiologies explain the association between hearing loss and cognitive decline. If incipient neurodegeneration influences hearing loss, genetic variants that increase risk of dementia should also increase risk of hearing loss (Figure 1).

**Figure 1.**
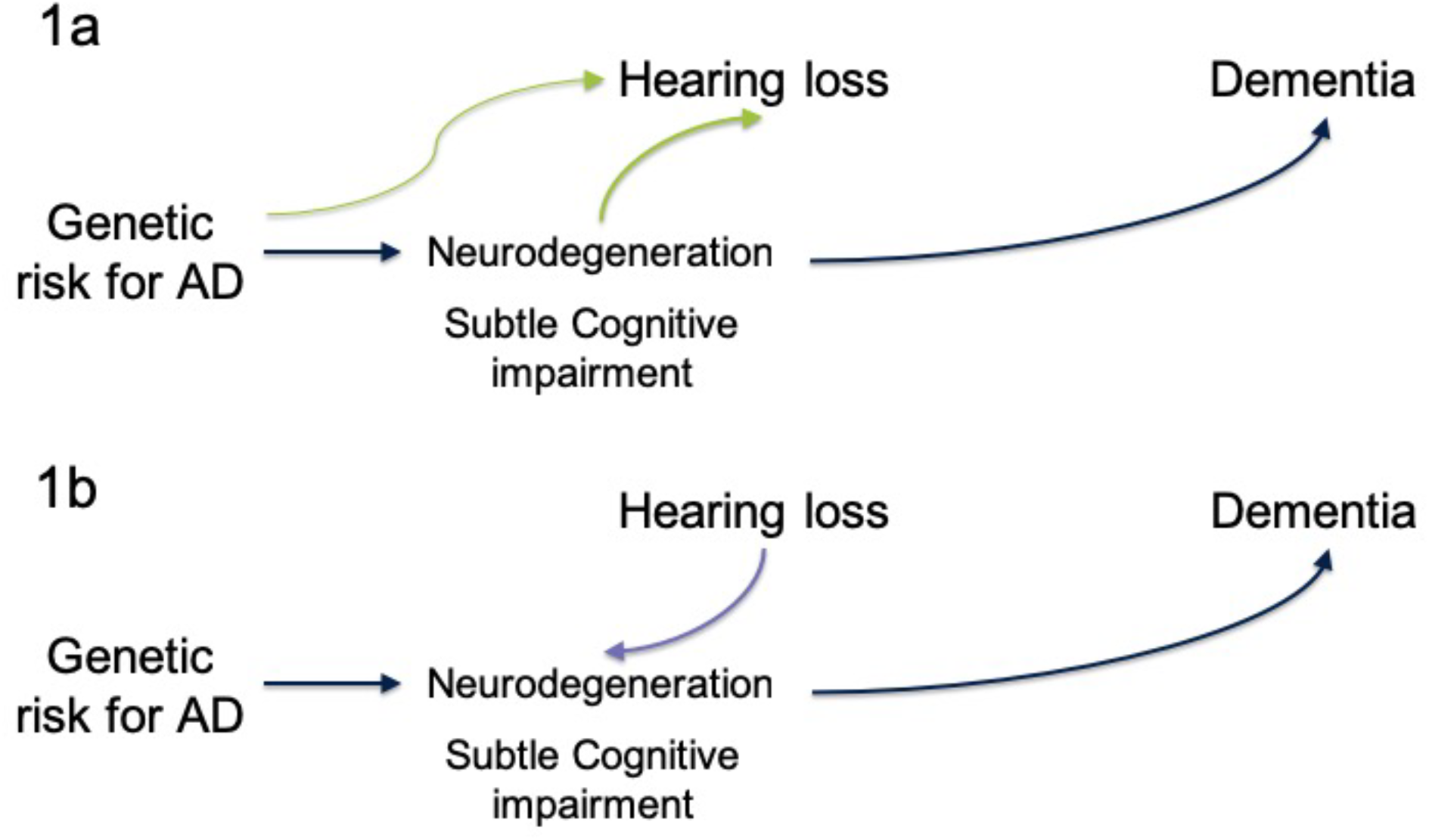
Conceptual models of AD genetic risk, dementia and hearing loss motivating analysis. Genetic variants known to increase AD risk can be used to distinguish between mechanisms that may explain the previously documented association between hearing loss and dementia risk. If incipient neurodegeneration or other shared etiologies influence hearing loss, then genetic variants that increase risk of dementia should also be associated with increased risk of hearing loss (1a). If hearing loss has effects on biological or social processes that increase risk of dementia, then genetic risk for AD should be independent of (not associated with) hearing loss (1b).

The objective of this study was to take advantage of genetic variation linked to AD, which is established at conception, to test the hypothesis that the link between hearing impairment and dementia may be explained by the influence of incipient AD on hearing loss. We used data for older adults participating in the UK Biobank and examined associations with both self-rated and an objective measure of hearing impairment.

## METHODS

### Study Setting and Participants

UK Biobank is an ongoing study of over 500,000 adults. Participants aged 40-69 years old were recruited 2006-2010 from across the UK to provide detailed information about themselves via computerized questionnaires, provide biologic samples, undergo clinical measurements, and have their health followed prospectively [24]. Ethical approval was obtained from the National Health Service National Research Ethics Service and all participants provided written informed consent.

The current analysis was restricted to UK Biobank participants age 56 or older (n=291,516) to focus on older adults where hearing loss was likely to be common. We excluded those with missing genetic information (n=8,560) or who were flagged as recommended for genetic analysis exclusion (n = 226), and those without any objective or subjective hearing assessments (n=693). We also excluded participants classified as of non-European genetic ancestry (n=37,122) based on a combination of self-report and genetic ancestry principal components, because genetic predictors of AD may differ by race/population stratification [25]. This left an analytic sample of 244,915 with at least one hearing measure. Objective hearing tests were available for 80,074 of these participants.

### Hearing

A Digit Triplet Test, a measure of speech-in-noise hearing ability, was incorporated into visits during 2009 and was completed by 80,136 eligible participants. Participants were asked to remove hearing aids prior to testing; those with cochlear implants were asked not to attempt the test. Participants were guided by a video demonstration and testing was performed via touchscreen and wore circumaural headphones (Sennheiser HD-25). The English speech materials for the UK Biobank Digit Triplet Test were developed at the University of Southampton and are described elsewhere [26,27]. Digit triplets (e.g. group of 3 monosyllabic digits such as 2-8-5) were presented in a total of 15 sets. A background noise matched to the spectrum of speech stimuli played simultaneously while digit triplets were presented. In initial triplets both noise and speech levels were adjusted together to a comfortable level. The speech level was then fixed, and noise levels increased or decreased adaptively after each triplet to estimate the signal-to-noise ratio at which a participant had 50% correct recognition of three digits. A Speech Reception Threshold (SRT) was used as the primary measure of objective hearing. SRT was calculated as the mean signal-to-noise ratio for triplets 8-15. Higher scores correspond to worse performance. In analyses, we used the SRT of the better hearing ear, following the approach of other studies [9,28]. Due to outliers and skewed distribution we log-transformed the SRT (calculated as log [SRT + 13] to account for negative SRT values) when analyzing it as a continuous measure of objective hearing. We also created a dichotomous indicator for poor objective hearing, defined as a SRT>-5.5 dB, corresponding with cut-offs used in previous research [9,28].

All participants were also asked about problems hearing (“Do you have any difficulty with your hearing?” and problem hearing in noise (e.g. “Do you find it difficult to follow a conversation if there is background noise (such as TV, radio, children playing). Possible answers were “Yes; “No”; “Do not know”; “Prefer not to answer”. We considered answers “Do not know” or “Prefer not to answer” as missing data.

### Genotyping and Genetic Risk Scores for AD

Genotyping of UK Biobank samples was conducted with two closely related arrays (Affymetrix using a bespoke BiLEVE Axiom array and Affymetrix UK Biobank Axiom array) and is described in detail elsewhere [29,30]. Briefly, all genetic data were quality controlled and imputed by UK Biobank [downloaded on 12/1/2017] to a reference panel that merged the 1,000 Genomes Phase 3 and UK10K reference panels. A secondary imputation was completed using the HRC reference panel and results from the HRC imputation were preferentially used at SNPs present in both panels. Before the release of the UK Biobank genetic data a stringent QC protocol was applied, which was performed at the Welcome Trust Centre for Human Genetics, and is described elsewhere [31]. We additionally excluded participants from the present study if they had non-European ancestry (as described above), were missing genetic information, or were recommended for genetic exclusion by UK Biobank due to high heterozygosity rate (after correcting for ancestry) or high missing rate [32].

We used summary results from the 2013 International Genomics of Alzheimer’s Project (IGAP) meta-analyzed genome-wide association study (GWAS) on late-onset AD in Caucasian populations[33] to calculate an AD genetic risk score (GRS) for each participant. The IGAP study identified 23 loci associated with AD, including 2 SNPs used to characterize APOE ε4 allele status. The GRS was based on the meta-analyzed β coefficients obtained in the IGAP’s stage 1 study, which included genotyped and imputed data (7,055,881 single nucleotide polymorphisms, 1000G phase 1 alpha imputation, Build 37, Assembly Hg19) of 17,008 Alzheimer’s disease cases and 37,154 controls. We calculated the GRS by multiplying each individual’s risk allele count for each locus by the β coefficient (expressed as the log OR) for that polymorphism (Table 1) and summing the products for all 23 loci. This step weights each SNP in proportion to its anticipated effect (either positive or negative) on AD risk. The scores can be interpreted as the log OR for AD conferred by that individual’s profile on the 23 SNPs compared to a person who had the major allele at each locus. With this construction, a one unit increase in the AD-GRS connotes a 2.7 times higher risk of AD. To assess the association with the AD-GRS beyond the effects of APOE ε4 (AD-GRS without APOE), we also calculated an alternative AD-GRS after removing the two SNPs associated with APOE. In a sensitivity analysis on the effects of APOE genotype alone, we created a score based on the two SNPs associated with APOE. From the APOE only scores we derived a dichotomous variable noting presence of at least one APOE ε4 allele.

**Table 1.** The 23 SNPs and their log odds ratio estimates for the Alzheimer’s disease genetic risk score (AD-GRS).

| **Marker Name** | **Chromosome** | **Closest gene** | **Effect Allele** | **Effect Estimate** |
| --- | --- | --- | --- | --- |
| rs6656401 | 1 | CR1 | A | 0.1567 |
| rs35349669 | 2 | INPP5D | T | 0.0663 |
| rs6733839 | 2 | BIN1 | T | 0.188 |
| rs190982 | 5 | MEF2C | G | -0.0799 |
| rs10948363 | 6 | CD2AP | G | 0.0978 |
| rs11771145 | 7 | EPHA1 | A | -0.1024 |
| rs1476679 | 7 | ZCWPW1 | C | -0.0783 |
| rs2718058 | 7 | NME8 | G | -0.0697 |
| rs28834970 | 8 | PTK2B | C | 0.0959 |
| rs9331896 | 8 | CLU | C | -0.1457 |
| rs10792832 | 11 | PICALM | A | -0.1297 |
| rs10838725 | 11 | CELF1 | C | 0.0753 |
| rs11218343 | 11 | SORL1 | C | -0.2697 |
| rs670139 | 11 | MS4A4E | T | 0.0803 |
| rs983392 | 11 | MS4A6A | G | -0.1084 |
| rs10498633 | 14 | SLC24A4-RIN3 | T | -0.1044 |
| rs17125944 | 14 | FERMT2 | C | 0.1223 |
| rs8093731 | 18 | DSG2 | T | -0.6136 |
| rs3865444 | 19 | CD33 | A | -0.0954 |
| rs4147929 | 19 | ABCA7 | A | 0.1348 |
| rs429358 | 19 | APOE | C | 1.3503 |
| rs7412 | 19 | APOE | T | -0.3871 |
| rs7274581 | 20 | CASS4 | C | -0.139 |

### Cognition

A measure of cognition was not necessary for our primary hypothesis test but was used to confirm previously established associations between the AD-GRS and cognition and between cognition and hearing impairment. The verbal reasoning assessment (also called fluid intelligence in some UK Biobank reports) was a visual touchscreen-based test that required participants to solve problems based on logic and reasoning. A summary score was calculated based on the number of correct responses to 13 questions within a two-minute time limit. Although other cognitive measures were assessed for some UK Biobank participants, we used this measure for several reasons: the verbal reasoning score was available for a much larger fraction of the eligible sample; it has an approximately normal distribution; it declines with age, and it is correlated with all other cognitive measures [34]. Associations between poor hearing and cognition using other cognitive tests have been reported previously [9].

### Other Covariates

Age and sex were reported during baseline assessment via touchscreen questionnaires. The UK Biobank provides principal components (PCs) related to genetic population stratification, we used the first 5 PCs in our analyses to adjust for population stratification.

### Statistical Analysis

We followed a modified Mendelian randomization analysis, which uses genetic variants as natural experiments to estimate effects of a risk factor of interest. This idea can be used to evaluate reverse causation as an explanation for observed associations, by finding genetic variants that influence an outcome, in our case, AD. Mendelian randomization approaches take advantage of the fact that genes have an established temporal order (i.e. determined at conception, prior to disease onset), and are therefore not susceptible to traditional confounders in observational studies to help improve casual inference in observational data [22]. If AD related genetic variants influence hearing loss, this cannot be attributed to the influence of hearing on dementia, but instead provides evidence for either reverse causation (from AD to hearing impairment) or a shared genetic etiology (Figure 1A). First, we confirmed that consistent with previous literature: 1) AD-GRS was associated with cognition (verbal reasoning) and 2) hearing impairment was associated with cognition in our sample. We tested our primary hypothesis by evaluating the association between AD-GRS and hearing impairment (both objective and self-reported measures). We estimated separate logistic regressions with AD-GRS as the predictor and each hearing impairment measure as the outcome (objective poor hearing in noise based on SRT, self-reported hearing problems, and self-reported problems with hearing in noise). A linear regression model tested the association between AD-GRS and continuous log SRT. All regression models included adjustment for age, sex, and 5 PCs to account for ancestry differences. Analyses were conducted in R (version 3.3.2). All tests were two-sided with α = 0.05 and we report 95% confidence intervals (CIs) to represent uncertainty in our effect estimates.

## RESULTS

Participant baseline characteristics are shown in Table 2. Objectively measure hearing speech-in-noise reception threshold (SRT) was on average −7.1 dB (Standard deviation [SD]: 1.8); 14% had poor objective hearing (SRT > −5.5 dB). Self-reported problems hearing were common (30-40%). 85% of those who reported any hearing problems also reported problems hearing in noise, 64.5% of those who reported any hearing problems in noise also reported problems hearing in general. There was less overlap between self-reported and objectively measured poor hearing (SRT > −5.5 dB); among those with poor objectively measured hearing, 52% had self-reported problems hearing and 57% had self-reported problems hearing in noise. However, objective hearing was significantly worse (higher value) in those with self-reported hearing problems (mean SRT was −6.7 dB [2.1] compared to −7.4 dB [SD]: 1.5 dB) and hearing problems in noise (mean SRT was −6.8 dB [2.1] compared to −7.4 dB [SD]: 1.5 dB) (both p<0.001, Kruskal Wallis test).

**Table 2.**
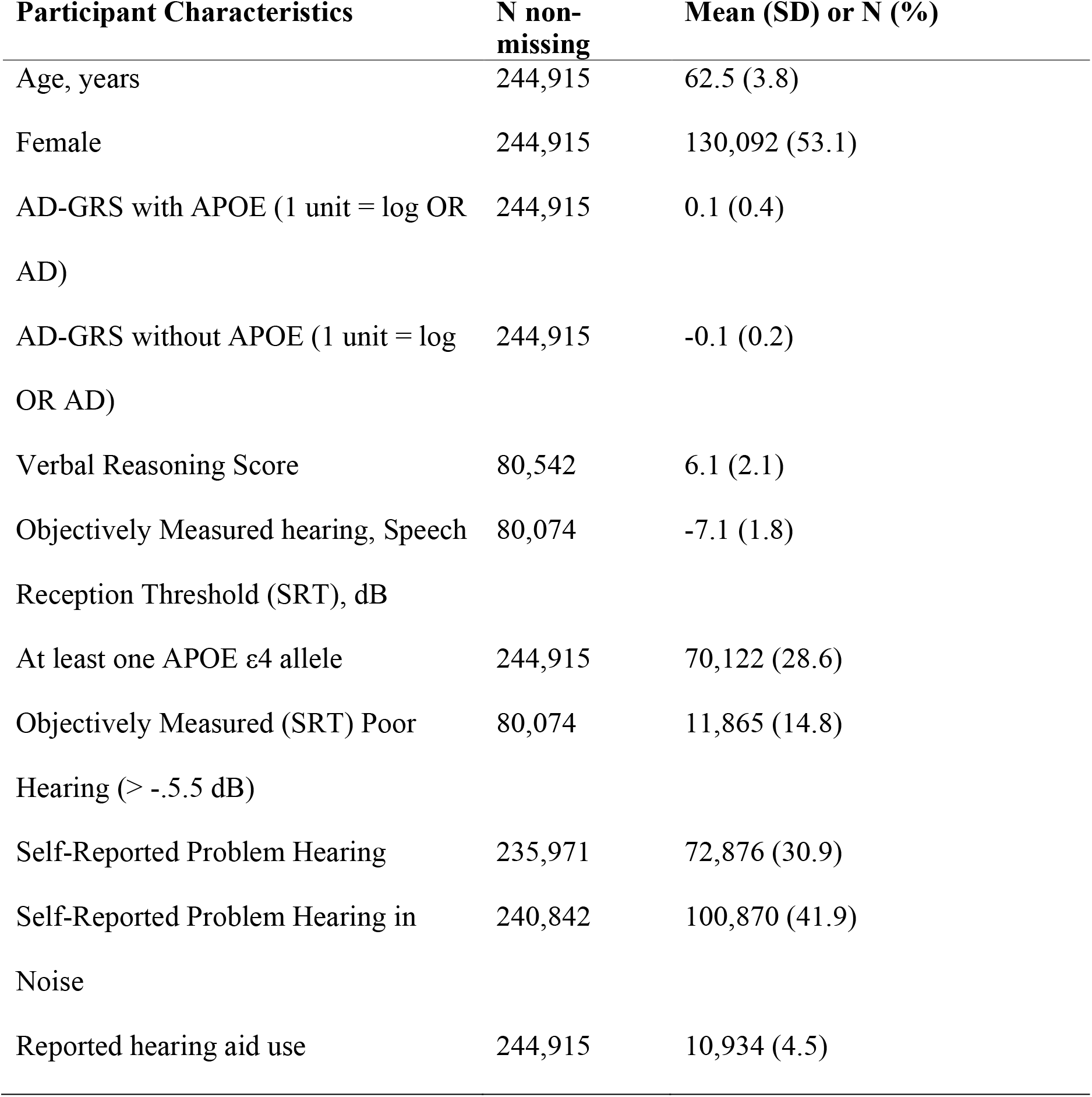
Characteristics of UK Biobank Participants Included in the Analyses

| **Participant Characteristics** | **N non-missing** | **Mean (SD) or N (%)** |
| --- | --- | --- |
| Age, years | 244,915 | 62.5 (3.8) |
| Female | 244,915 | 130,092 (53.1) |
| AD-GRS with APOE (1 unit = log OR AD) | 244,915 | 0.1 (0.4) |
| AD-GRS without APOE (1 unit = log OR AD) | 244,915 | -0.1 (0.2) |
| Verbal Reasoning Score | 80,542 | 6.1 (2.1) |
| Objectively Measured hearing, Speech Reception Threshold (SRT), dB | 80,074 | -7.1 (1.8) |
| At least one APOE ε4 allele | 244,915 | 70,122 (28.6) |
| Objectively Measured (SRT) Poor Hearing (> -5.5 dB) | 80,074 | 11,865 (14.8) |
| Self-Reported Problem Hearing | 235,971 | 72,876 (30.9) |
| Self-Reported Problem Hearing in Noise | 240,842 | 100,870 (41.9) |
| Reported hearing aid use | 244,915 | 10,934 (4.5) |

### Confirming Associations of Hearing and AD GRS with Cognition

Every measure of poor hearing was associated with worse verbal reasoning in the subset of participants with verbal reasoning measures (Table 3). Higher AD-GRS with APOE was associated with worse verbal reasoning (Table 3), however, the AD-GRS without APOE was not significantly associated with verbal reasoning.

**Table 3.** Linear regression coefficients predicting verbal reasoning scores as a function of self-reported and objective measures of hearing and AD-GRS

| Predictors | Verbal Reasoning Score<br>$\beta$ (95% CI) |
| --- | --- |
| *Objective Hearing Measures* | N=78,010 |
| Poor SRT | -1.28 (-1.54, -1.03) |
| SRT (continuous) | -0.13 (-0.14, -0.12) |
| *Self-Reported Hearing Measures\** |  |
| Problem hearing (n=76,156) | -0.08 (-0.12, -0.05) |
| Problem hearing in noise (n=78,642) | -0.08 (-0.11, -0.05) |
| *Alzheimer's Disease Genetic Risk Scores\*\** | N= 80,542 |
| AD-GRS with APOE | -0.04 (-0.07, -0.0002) |
| AD-GRS without APOE | 0.02 (-0.07, 0.10) |

\*Adjusted for age, sex
\*\* additionally adjusted for 5 PCs to account for population stratification
Abbreviations: AD-GRS, Alzheimer's disease genetic risk score; SRT, Speech Reception Threshold

### AD GRS and Hearing Associations

AD-GRS with and without APOE was associated with objectively measured poor hearing (SRT) (p<0.02 for both GRS) and self-reported problems hearing in noise (p<0.01 for both GRS) (Table 4). The association of the AD-GRS and self-reported problems hearing was in the same direction but not statistically significantly (p=0.05 for AD-GRS and p=0.08 for AD-GRS without APOE). Effect sizes across all hearing outcomes ranged from a 2% to 13% increased odds of poor hearing per 1 unit increase in AD-GRS; to put this into more interpretable terms, an increase in the AD-GRS that would roughly triple the odds of AD would also increase risk of hearing impairment by 2-13%. There was a trend towards slightly higher effect sizes in analyses using objectively measured poor hearing. The AD-GRS with APOE (p=0.03) (but not AD-GRS without APOE; p=0.3) was associated with objectively measured SRT on the continuous scale (Figure 2 shows effects by sample quartiles of AD-GRS with APOE). An GRS based on APOE alone was not significantly associated with poor SRT (OR =1.04; 95% CI: 0.98, 1.10; p=0.2).

**Table 4.** Odds Ratios for the Association between AD-GRS and Three Measures of Hearing

| **AD GRS** | **Objectively Measured Poor Hearing SRT OR (95% CI)** | **Self-Reported Problems Hearing OR (95% CI)** | **Self-Reported Problems Hearing in Noise OR (95% CI)** |
| --- | --- | --- | --- |
| N | 80,074 | 235,971 | 240,842 |
| AD-GRS with APOE | 1.06 (1.01, 1.11) | 1.02 (1.00, 1.05) | 1.03 (1.005, 1.05) |
| AD-GRS without APOE | 1.13 (1.01, 1.27) | 1.05 (0.99, 1.10) | 1.07 (1.02, 1.13) |

Abbreviations: AD-GRS, Alzheimer's disease genetic risk score; SRT, Speech Reception Threshold

**Figure 2.**
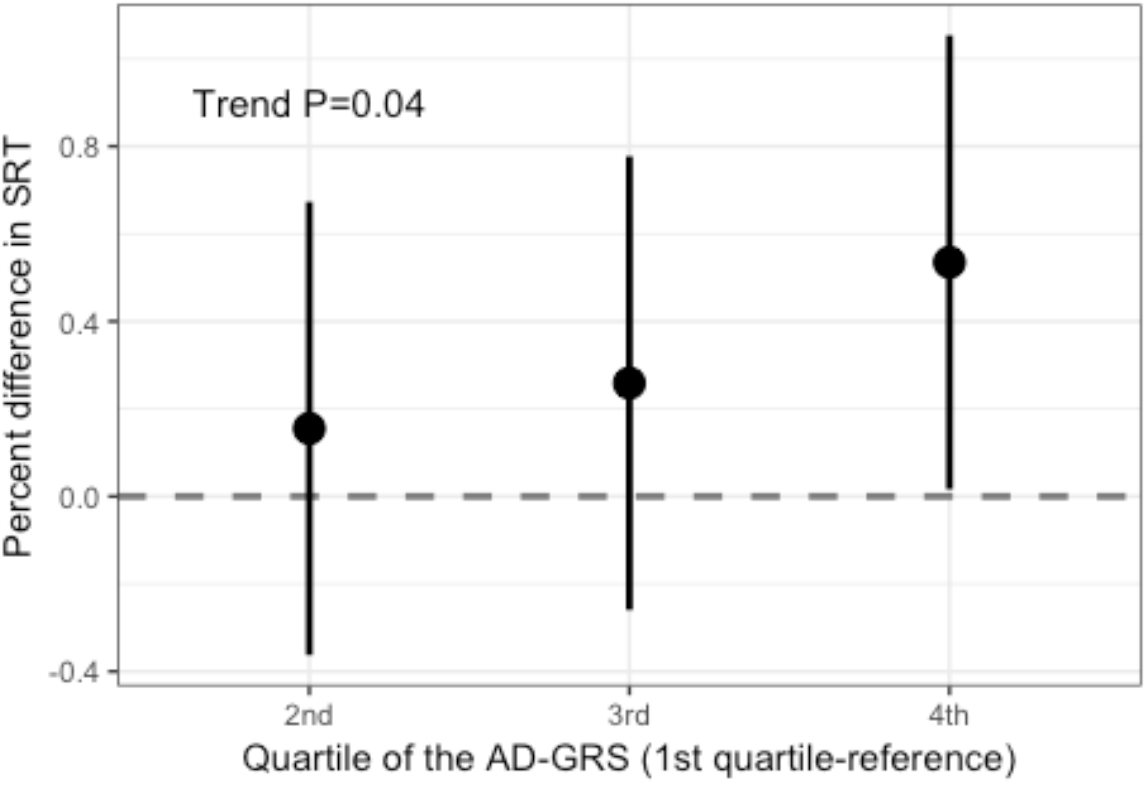
Association between objectively measured hearing (Speech Reception Threshold [SRT]) and quartiles of Alzheimer’s disease genetic risk score (AD-GRS) with APOE (n=80,074). There was a significant trend (p=0.04) for increasing quartile of AD-GRS and worse SRT (positive change), however, only the 4^th^quartile was significantly worse SRT than the 1^st^ quartile (reference group). Based on a linear regression of log SRT adjusted for age, sex, and 5 PCs to account for population stratification

## DISCUSSION

In this study, we found that higher AD-GRS was associated with worse hearing in a large sample of middle-aged and older adults. Findings were consistent across self-reported and objective hearing measurements; the largest effect estimates were for the association of AD-GRS and objectively measured SRT. Results were similar using either the AD-GRS with APOE or the AD-GRS without APOE. Although effect estimates were small, these findings provide evidence that genetic risk for AD is associated with worse hearing, particularly speech reception in noise. This evidence supports the hypothesis that underlying dementia disease processes or shared etiologies influence hearing ability and at least partially explain the association between hearing loss and cognition.

Prior studies have found associations of hearing with cognition[4–7] and structural brain volumes [35,36], however, the mechanisms underlying this association remain unclear. We used polygenetic risk scores for AD to help address a key limitation of traditional observational approaches to evaluating the direction of the association linking poor hearing and dementia. Given this sample was relatively young and healthy at enrollment, participants with high AD-GRS are disproportionately likely to be experiencing early effects of AD. Following the AD pathophysiologic cascade [37], neurodegeneration may occur many years prior to development of dementia symptoms. Additionally, very subtle cognitive impairments can be detected many years prior to dementia diagnosis. These changes may in turn lead to worse hearing, consistent with our finding that higher genetic AD risk is associated with worse hearing. Our findings may reflect changes in central hearing rather than peripheral hearing since detecting speech in noise requires central hearing processing and general cognitive abilities to discriminate digits. Although peripheral hearing loss is still the primary predictor of speech reception in noise [38], future research should examine genetic risk and audiometric measures.

Another interpretation of our findings is that the genetic variants related to AD have pleiotropic pathways via which they influence hearing. The genetic risks may also influence non-neurodegenerative factors, such as vascular disease, that may act as shared risk factors for hearing loss and cognitive decline. Several AD genes, including APOE ε4 allele impact lipid metabolism [39] and APOE genotype is associated with cardiovascular risk factors as well [40]. However, a genome-wide association study of neuropathologic contributors to dementia found that AD genes are associated with AD neuropathology, including amyloid plaque and neurofibrillary tangle burden, but not with vascular brain injury [41]. Furthermore, AD genes have not been linked as primary predictors of impaired hearing, in fact few genes are significantly associated with age-related hearing loss [42]. This lends further evidence to the hypothesis that the AD disease process may contribute to impairment in hearing but does not exclude the possibility of pleiotropic effects. Furthermore, our analyses cannot distinguish whether these findings are because of neurodegeneration in auditory brain regions or cognitive impairment. Regardless of this ambiguity in interpretation, our results suggest that the association between hearing impairment and dementia risk in conventional observational studies partially reflects a shared pathway associated with the AD-GRS.

Our findings do not exclude the possibility that hearing loss causes cognitive decline, as multiple mechanisms may contribute to the relationship between hearing loss and dementia. The Lancet commission on dementia prevention calculated a relative risk of impaired hearing on dementia of 1.9 [11], equating to an odds ratio (OR) of 2.0-2.5, depending on the prevalence of dementia. This estimated association between hearing and dementia is much higher than could be attributed to the effects of known AD genetic risk on impaired hearing in this study ORs= 1.05-1.13. This suggests other mechanisms link hearing and dementia but does not establish that those other mechanisms are necessarily a causal effect of hearing on AD. The estimates in our current analysis are specific to genetically determined AD and do not capture pleiotropic effects of unmeasured determinants of AD, which may underestimate the effect of AD genetic risk on hearing. Interventions to improve hearing may not be as effective at reducing dementia burden if the association between hearing and dementia is partly explained by underlying AD process or shared etiologies. Future studies will be needed to determine whether hearing loss accelerates cognitive decline independent of shared etiologies.

There are several important caveats and limitations to our analysis. Our results are potentially influenced by selection bias due to selective survival as the AD-GRS is associated with mortality [43]. However, this effect is likely limited because the sample is relatively young and healthy with a low mortality rate, especially due to dementia. Genetic risk for AD only explains a small percentage of variation in cognitive impairment and AD diagnosis, thus there is substantial variation in risk for AD that is not captured by our risk score. This variation reduces the ability to detect associations, suggesting the strength of the association between dementia processes and hearing is even higher than we found. However, there are considerable strengths to this study, particularly the size of sample, presence of self-rated and an objective measure of hearing function, and innovative use of a modified Mendelian randomization approach to test whether early-stage dementia processes may influence hearing function. By using genetic risk to address this question, we circumvent the central challenge in interpreting prior results showing that hearing loss and dementia risk are associated.

We used variation in genetic risk for AD as an approach to test whether the dementia disease processes influence hearing (e.g. reverse causation). In support of this hypothesis, we found that higher genetic risk for AD was positively associated with worse hearing ratings and worse speech in noise testing. Our findings suggest underlying shared etiology may help explain the link between poor functional hearing and dementia. Individuals with poor hearing may be more likely to have preclinical AD than those with normal hearing. Additional research will be needed to replicate findings in other samples and with measures of peripheral hearing loss as well as to explicitly test whether hearing loss accelerates cognitive decline.

## ACKNOWLEDGEMENTS

We thank UK Biobank participants and staff. The UK Biobank project is funded by the Medical Research Council, The Wellcome Trust, Department of Health for England and Wales, North West Regional Development Agency and the Scottish Executive.

This work was supported by NIA grants T32-AG-049663 (WDB), K24-AG031155 (KY), and RF1AG052132-01 (RAW and MMG).

